# Freshwater *Chlorobia* exhibit metabolic specialization among cosmopolitan and endemic populations

**DOI:** 10.1101/2020.09.10.291559

**Authors:** Sarahi L. Garcia, Maliheh Mehrshad, Moritz Buck, Jackson M. Tsuji, Josh D. Neufeld, Katherine D. McMahon, Stefan Bertilsson, Chris Greening, Sari Peura

## Abstract

Photosynthetic bacteria from the class *Chlorobia* (formerly phylum *Chlorobi*) sustain carbon fixation in anoxic water columns. They harvest light at extremely low intensities and use various inorganic electron donors to fix carbon dioxide into biomass. Until now, most information on their functional ecology and local adaptations came from isolates and merely 26 sequenced genomes that are poor representatives of natural populations. To address these limitations, we analyzed global metagenomes to profile planktonic *Chlorobia* cells from the oxyclines of 42 freshwater bodies, spanning subarctic to tropical regions and encompassing all four seasons. We assembled and compiled over 500 genomes, including metagenome-assembled genomes (MAGs), single-cell genomes (SAGs), and reference genomes from cultures, clustering them into 71 metagenomic operational taxonomic units (mOTUs) or “species”. Of the 71 mOTUs, 57 were classified as genus *Chlorobium* and these mOTUs varied in relative abundance up to ~60% of the microbial communities in the sampled anoxic waters. Several *Chlorobium*-associated mOTUs were globally distributed whereas others were endemic to individual lakes. Although most clades encoded the ability to oxidize hydrogen, many were lacking genes for the oxidation of specific sulfur and iron substrates. Surprisingly, one globally distributed Scandinavian *Chlorobium* clade encoded the ability to oxidize hydrogen, sulfur, and iron, suggesting that metabolic versatility facilitated such widespread colonization. Overall, these findings provide new insights into the biogeography of the *Chlorobia* and the metabolic traits that facilitate niche specialization within lake ecosystems.

**Importance:** The reconstruction of genomes from metagenomes has enabled unprecedented insights into the ecology and evolution of environmental microbiomes. We applied this powerful approach to 274 metagenomes collected from diverse freshwater habitats that spanned oxic and anoxic zones, sampling seasons, and latitudes. We demonstrate widespread and abundant distributions of planktonic *Chlorobia*-associated bacteria in hypolimnetic waters of stratified freshwater ecosystems and pinpoint nutrients that likely fuel their electron chains. Being photoautotrophs, these *Chlorobia* organisms also have the potential to serve as carbon sources that support metalimnetic and hypolimnetic food webs.

## Introduction

Although oxygenic phototrophs dominate contemporary carbon fixation, anoxygenic phototrophs have been important over planetary timescales and continue to occupy important niches in a wide range of ecosystems (1). Among the anoxygenic phototrophs, members of the class *Chlorobia* (green sulfur bacteria; formerly phylum *Chlorobi*) are physiologically remarkable. Known *Chlorobia* members are obligate photolithoautotrophic anaerobes (2) that leverage bacteriochlorophyll-rich organelles called chlorosomes to harvest light at extremely low intensities (i.e., in the range of 1–10 nmol photons m^−2^ s^−1^) (3). Such efficient light harvesting determines the ecological niche of these bacteria at the lowermost strata of the photic zone in stratified water columns, where the least amount of light is available (4, 5). Studies on isolates obtained from different aquatic systems and microbial mats have revealed that members of class *Chlorobia* use electrons derived from reduced sulfur compounds and/or hydrogen to fix carbon dioxide via the reverse tricarboxylic acid (TCA) cycle (6–8). Although several strains can oxidize thiosulfate, they almost universally use sulfide as electron donor for CO_2_ reduction and source of sulfur for assimilation (2). Several members of the class *Chlorobia* are additionally known to oxidize ferrous iron (Fe^2+^) in a process called photoferrotrophy (9). Such photoferrotrophs include *Chlorobium ferrooxidans*, which obtains sulfur via assimilatory sulfate reduction, along with three other recently characterized strains (10–13). Although all characterized *Chlorobia* pure cultures can grow with carbon dioxide (CO_2_) or bicarbonate as their sole carbon source, many isolates have been reported to assimilate simple organic acids, such as acetate and pyruvate, under photomixotrophic or photoheterotrophic conditions (2). Moreover, most *Chlorobia* cells also encode the potential for nitrogen fixation (14–16). Together, this metabolic capacity and versatility enables members of the class *Chlorobia* to colonize dimly lit anoxic environments.

In stratified freshwater ecosystems, *Chlorobia* microorganisms have particularly important ecological and biogeochemical roles (17–20). In such systems they accumulate to form dense populations at certain strata, manifesting as deep chlorophyll maxima (21), and they also play prominent roles in carbon budgets of such lakes (22, 23) while concomitantly contributing to various processes in sulfur, iron, hydrogen, and nitrogen cycling (16, 24, 25). However, our knowledge of the geographical distribution and metabolic capabilities of *Chlorobia* members within natural planktonic communities remains incomplete. This is because most of our understanding of the metabolic capabilities and ecophysiological strategies of this class come from studies of approximately 100 isolates (4), with genomes available for 26 of these representatives (2, 26, 27), and cultures are often poor representatives of natural populations. The diversity and distributions of planktonic *Chlorobia* members are largely unknown. Moreover, the question still remains: how are genes conferring distinct ecophysiological traits distributed among *Chlorobia*-associated genomes from geographically distinct water bodies?

In this study, we used a cultivation-independent approach to gain a more comprehensive understanding of the natural distributions and metabolic capabilities of *Chlorobia* members within 42 freshwater bodies that are distributed across boreal, subarctic, and tropical regions in Europe and North America (Figure 1). Through this extensive study we assembled and analyzed over 500 genomes of *Chlorobia* members, including 465 metagenome-assembled genomes (MAGs) and 21 single-amplified genomes (SAGs). These were further clustered into 71 metagenome operational taxonomic units (mOTUs; where each mOTU is a cluster of genomes that share 95% average nucleotide identity). We observed that some of these mOTUs exhibited cosmopolitan distributions, whereas other lineages were locally constrained and possibly adapted to specific lakes or regions. In addition, clades varied in their encoded metabolic capabilities and we found several mOTUs that appear consistent with substrate specialization. These findings reveal further metabolic versatility and niche specialization within the abundant members of *Chlorobium*-associated bacteria.

**Figure 1.**
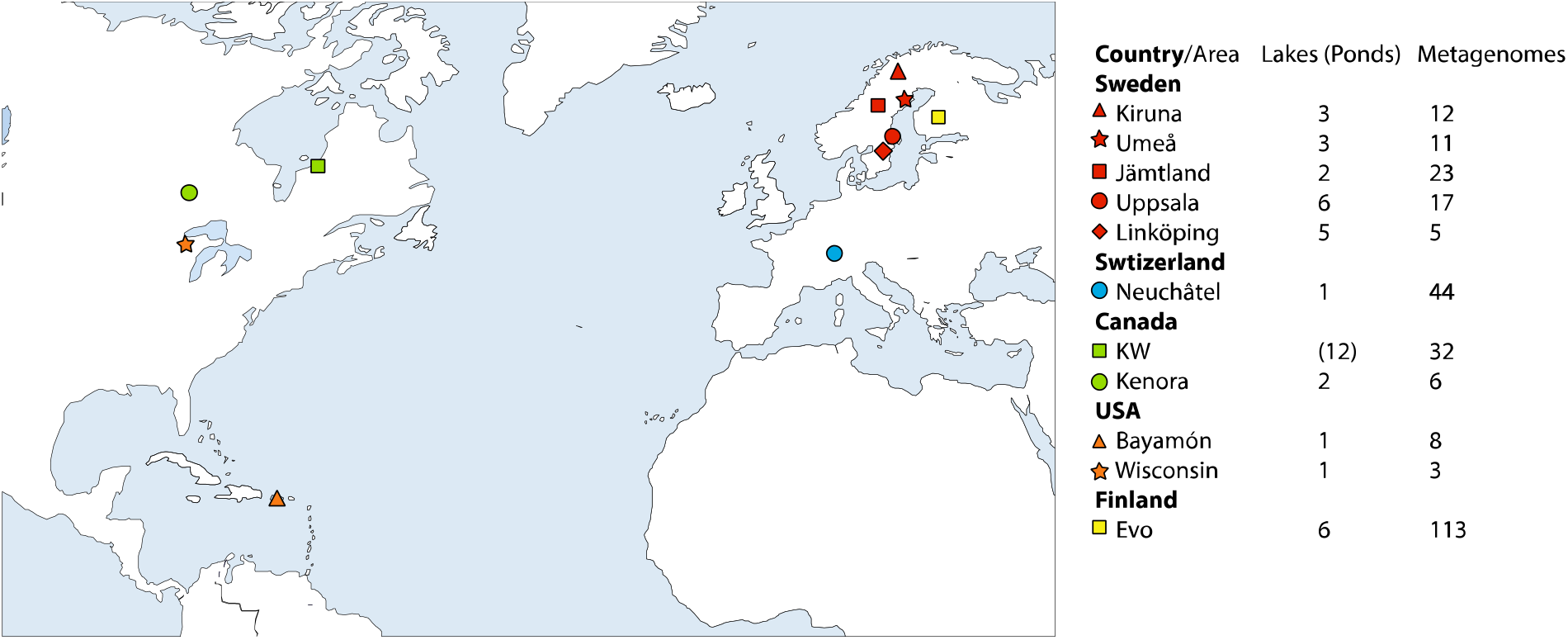
Areas from which we gathered lake or pond metagenomes for mapping and recovery of metagenome-assembled genomes (MAGs) and single-cell genomes (SAGs). Symbols are colored by country, and a unique shape is used for each area within the same country. KW stands for Kujjuarapik-Whapmagoostui.

## Results and discussion

### Metagenome assembly and characterization reveals that most planktonic members of the class Chlorobia affiliate with the genus Chlorobium

In this study we investigated the planktonic *Chlorobia*-affiliated populations of 30 lakes and 12 ponds in different regions of Europe and North America (Figure 1). Most of these samples were collected with the objective of investigating depth-associated changes in taxonomic diversity in different water masses that capture oxic epilimnion, metalimnion, and anoxic hypolimnion conditions (28). Following assembly and binning of individual metagenomes, we compiled 454 new metagenome-assembled genomes (MAGs) that affiliated with the class *Chlorobia*. Moreover, we collected 19 SAGs belonging to the class *Chlorobia* from two of the sampled lakes. To characterize the phylogenetic distributions of these new MAGs and SAGs, we further compiled 25 genomes available from the Genome Taxonomy Database (GTDB) and 11 MAGs available from previous studies (26, 29–31). In total, the dataset included 509 genomes including the MAGs, SAGs, and complete genomes of isolates (Table S1). Genome completeness varied from 50 to 100%, with an average of 89.0% and a median of 94.5% (Table S1). Contamination was below 5% for all genomes according to CheckM (32).

We clustered these genomes by 95% average nucleotide identity (ANI). This clustering into mOTUs enabled us to gain a more comprehensive understanding of the genomic structure of genus *Chlorobium* members. By clustering of >500 genomes at a 95% ANI threshold, previously shown to unite classical species definitions and separate sequenced strains into consistent and distinct groups (33–35), we obtained 71 metagenomic operational taxonomic units (mOTUs) or “species” belonging to the class *Chlorobia* (Table S2). Of those, 57 mOTUs were classified as members of the genus *Chlorobium* under the GTDB taxonomy (Figure 2 and table S2) (26). These 57 mOTU genomes had an average completeness of approximately 90%. Their mean estimated length was 2.6 Mbp, with the lowest estimated size being 2.1 Mbp and the highest 3.7 Mbp, which is consistent with previous studies showing that *Chlorobia* genomes from isolates range in size from 1.9 to 3.3 Mbp (2). Among the 57 *Chlorobium*-associated mOTUs, 12 were composed exclusively of genomes from previously described isolates (2), including *Chlorobium (Chl.) phaeobacteroides, Chl. limicola, Chl. luteolum, Chl. ferrooxidans and Chl. phaeoclathratiforme* (Figure 2). None of the mOTUs included both isolate genomes and environmental MAGs. Whereas several of the cultured members of the class *Chlorobia* were isolated from lakes (36), our analysis suggests that the majority of natural diversity within freshwater planktonic members of the class *Chlorobia* are found within uncultivated members of the genus *Chlorobium*.

**Figure 2.**
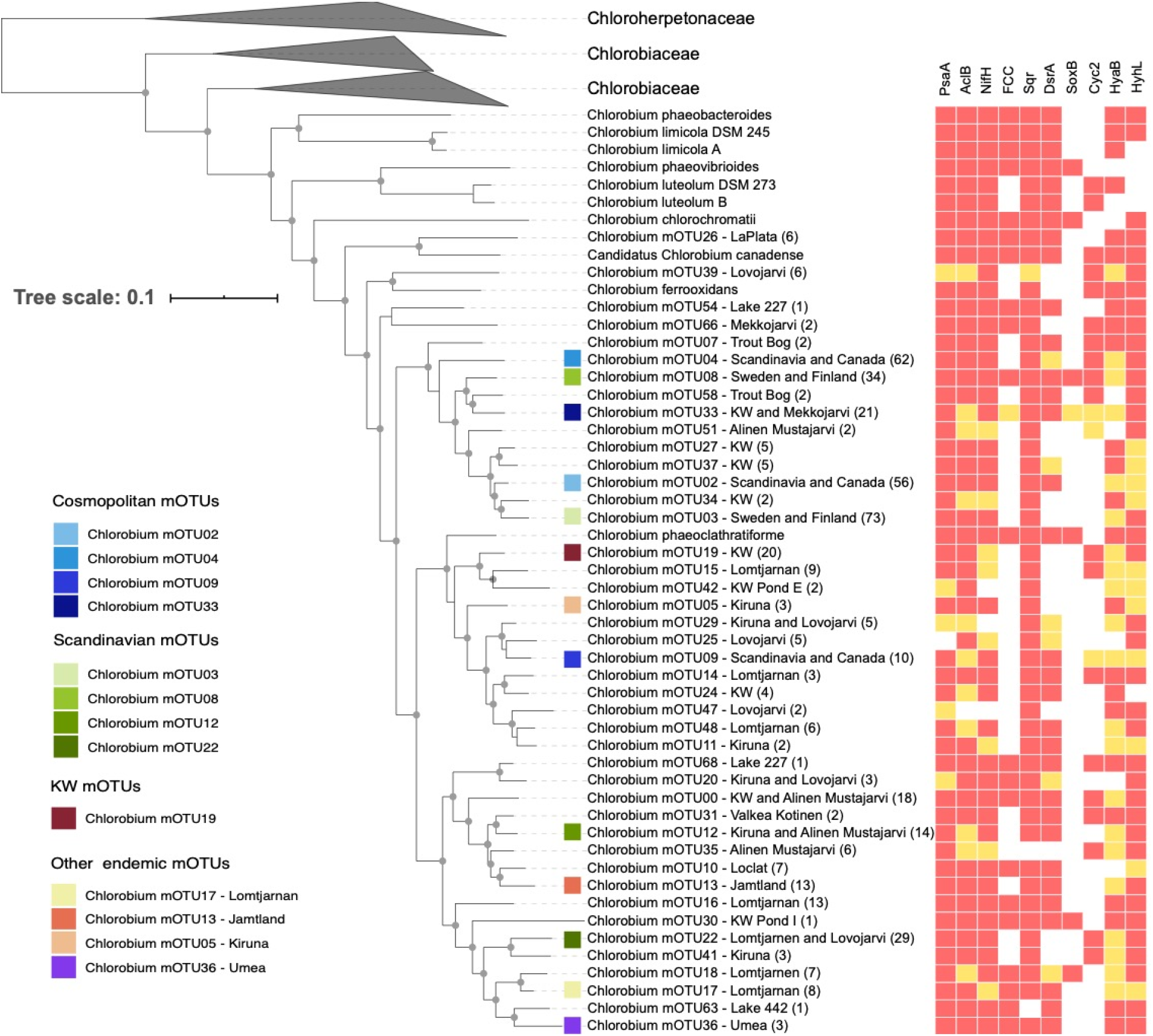
Phylogenetic relationship and metabolic capabilities of Chlorobium shows cosmopolitan and endemic mOTUs as non-monophyletic. The tree was constructed using GTDB-Tk “de-novo” alignment, and it was annotated and curated in iTOL. Isolates are shown in the tree with species names instead of mOTU number. Other tips on the tree show the genus name and mOTU number followed by location where the genomes were assembled from. If the MAGs where assembled from more than one lake in the same area, then the name of the area is written. If the MAGs where assembled from more than one area in the same country, then the name of the country is written. Lake La Plata is in Bayamon, Lakes 227 and 442 are in the IISD-ELA (near Kenora, Canada), Lake Lovojarvi, Lake Mekkojarvi, Lake Valkea Kotinen and Lake Alinen Mustajarvi are in Evo, Lake Trout Bog is in Wisconsin, Lake Loclat is in Neuchatel and Lake Lomtjarnen is in Jamtland. Numbers in parenthesis shows the number of genomes in each mOTU. The Genus *Chlorobium* is part of the Family *Chlorobiaceae*. Other genomes in different Genera within the family *Chlorobiaceae* are clustered. The tree also includes information about the presence/absence of some gene products (PsaA – Photosystem I P700 chlorophyll a apoprotein A1, AcIB – ATP-citrate lyase beta-subunit, NifH – nitrogenase iron protein, FCC – flavocytochrome c sulfide dehydrogenase, Sqr – sulfide-quinone oxidoreductase, DsrA – reverse dissimilatory sulfite reductase, SoxB – thiosulfohydrolase, Cyc2 – iron-oxidizing outer-membrane c-type cytochrome, HyaB – group 1d [NiFe]-hydrogenase large subunit, HyhL – group 3b [NiFe]-hydrogenase). In the heatmap red means the gene is present in the core genome of the mOTU, and yellow means it is present in the accessory genome of the mOTU, and white means the gene is absent in the mOTU, see the methods for details. The 13 most abundant *Chlorobium* mOTUs in the investigated environments have a colored square before the name of the mOTU. Gray circles in the tree represent bootstrap values higher that 50%.

### The genus Chlorobium includes both cosmopolitan and endemic lineages

By identifying MAGs from different source locations that cluster within the same mOTUs, we were able to determine whether a “species” is widely distributed and putatively cosmopolitan, or in contrast endemic to specific locations. To investigate the abundance and prevalence of each reconstructed mOTU within resident *Chlorobia* assemblages, we mapped metagenomic reads from the lake and pond datasets (Table S3) against the genome collection (Table S1). To reduce noise, we mapped reads with 100% identity to the reference and applied a cutoff for presence below 0.03% metagenomic read abundance. Considering these cutoffs, reads of our lake/pond dataset mapped to 45 (out of 71) *Chlorobia* mOTUs, of which 42 mOTUs classified as members of the genus *Chlorobium* (Figure 3). None of these mOTUs contained any of the cultivated *Chlorobium* representatives. This provides further evidence that uncultivated *Chlorobium* members are the dominant members of the planktonic *Chlorobia* present in sampled lakes and ponds.

**Figure 3.**
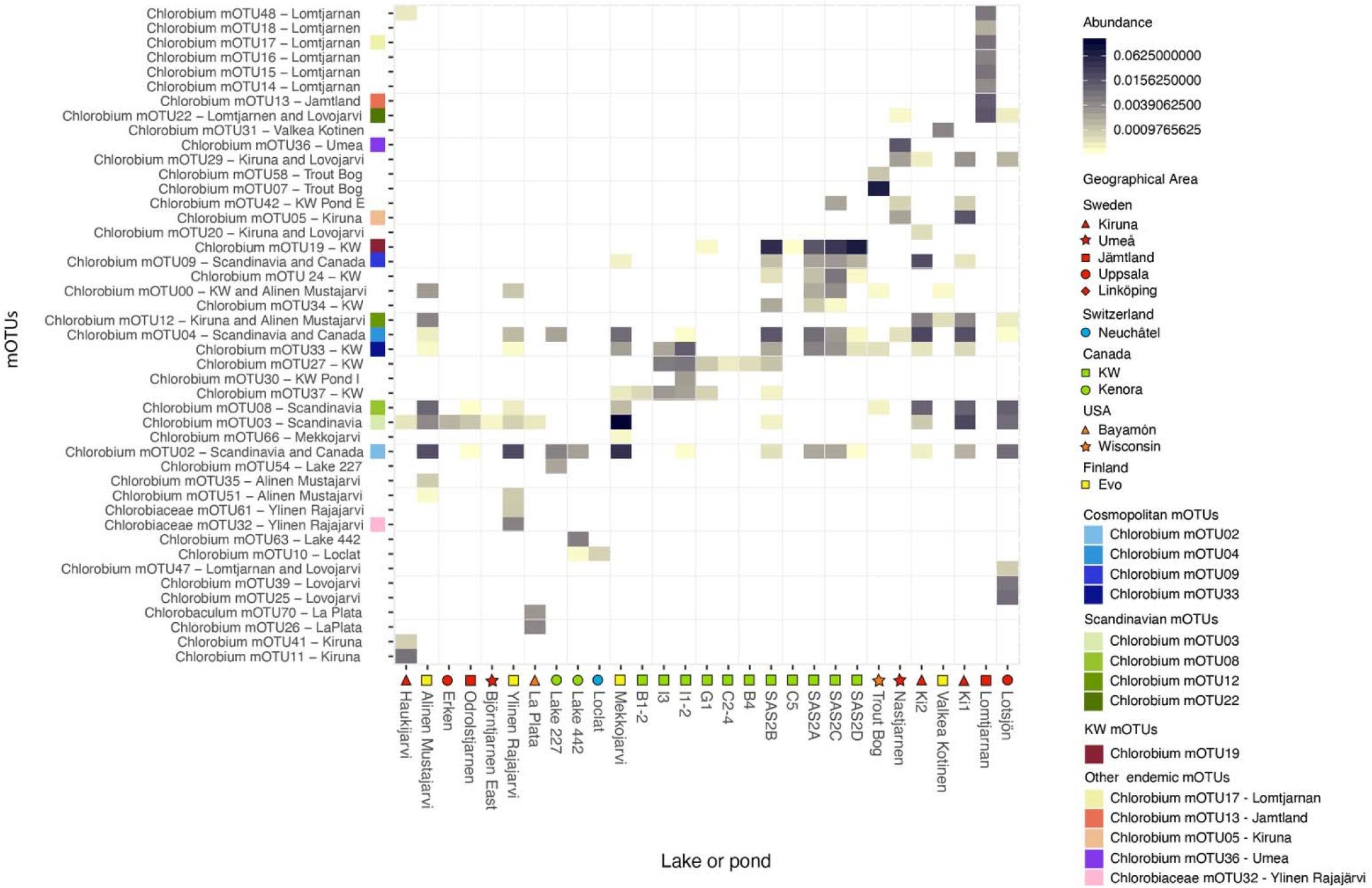
Relative abundance of the metagenome reads of the 45 abundant *Chlorobia* mOTUs in lake and pond metagenomes shows some mOTUs are truly endemic while others are more widespread or cosmopolitan. Top 14 *Chlorobia* mOTUs are color coded according to the location they were assembled from. Metagenomes were subsampled to one million reads. Reads were then mapped competitively with 100% identity cutoff to all genomes in all mOTUs using BBmap. The reads were normalized to relative abundance of reads per metagenome. Relative abundances from all depths and timepoints in a given lake/pond were averaged. The cutoff for presence of a mOTU to be included in the figure is 0.03% read abundance per lake/pond. The name of the mOTUs includes geographical information about the origin of the genomes as in Figure 2. The lakes or ponds in the x axis include a symbol for the region where they are located.

Out of the 42 locations represented by this study, we found that metagenomes sequenced from 18 lakes (of 30 total) and 11 ponds (of 12 total) contained *Chlorobia* reads (Figure 3). However, for 10 of the lakes that did not show *Chlorobia* read abundance, we only have epilimnion samples with full oxygenation, and so it was expected to not find *Chlorobia* reads or genomes. If we deduct these 10 lakes, we found *Chlorobia* reads in most of the metagenomes from anoxic freshwaters. In our data, we found that four mOTUs with MAGs reconstructed from assemblies originating on both studied continents (i.e., mOTU02, mOTU04, mOTU09, and mOTU33; in blue, Figures 2, 3, 4, and 5) and so define them as cosmopolitan because of their distribution pattern. Moreover, metagenomic read recruitment confirmed that these cosmopolitan mOTUs were present in lakes and ponds from both studied continents (Figure 3). In contrast, the other 41 mOTUs were less widely distributed and were found either in just one lake, in one geographical area, or in Scandinavia. The apparent cosmopolitan or locally constrained clades did not appear to be monophyletic, but rather distributed across the phylogenetic tree of the class *Chlorobia* (Figure 2).

**Figure 4.**
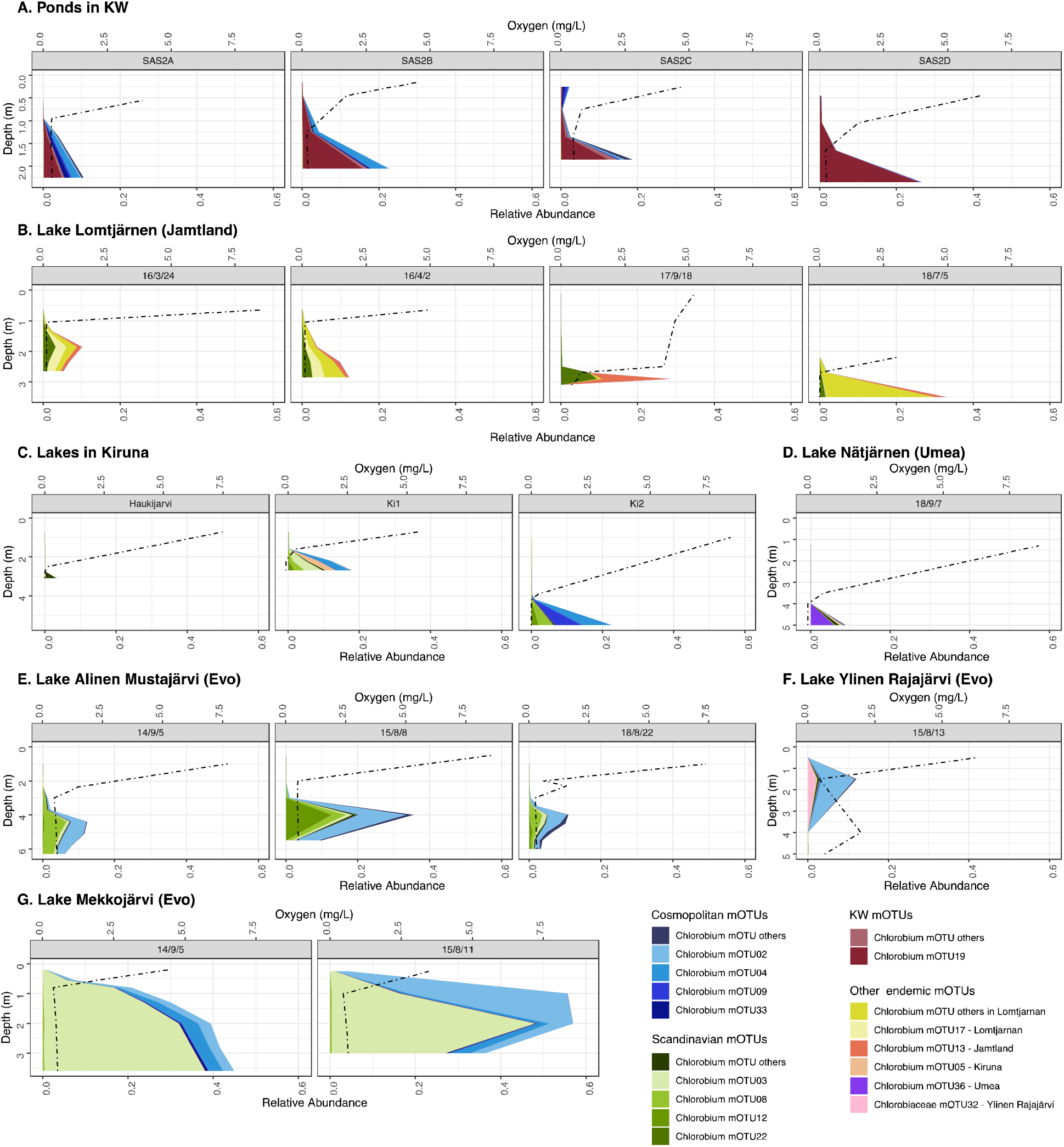
Distribution of environmentally abundant mOTUs across depth profiles in the lake and pond datasets with depth discrete sampling and high Chlorobium abundances shows abundance of *Chlorobia* below the oxycline. Note that the 14 most abundant *Chlorobia* mOTUs have individual color coding and while the remaining abundant mOTUs are combined in a category labeled “other”. For cosmopolitans, this include mOTU 00 and 42 and for Scandinavians this include mOTU 35, 29, 11, 31, 51 and 66. Other mOTUs in Kuujjuarapik-Whapmagoostui include mOTU 37, 24, 27, 34 and 30. Finally other mOTUs in the category endemic include mOTU 14, 15, 16, 18 and 48. Oxygen is represented by a dashed line. (A) Ponds in the KW area. (B) Different time points for Lake Lomtjärnen in Sweden. (C) Different lakes in the Kiruna area. (D) Lake Nästjärnen in Sweden. (E) Different time points of Lake Alinen Mustajärvi in Finland. (F) Lake Ylinen Rajajärvi. (G) Different time points for Lake Mekkojärvi in Finland.

**Figure 5.**
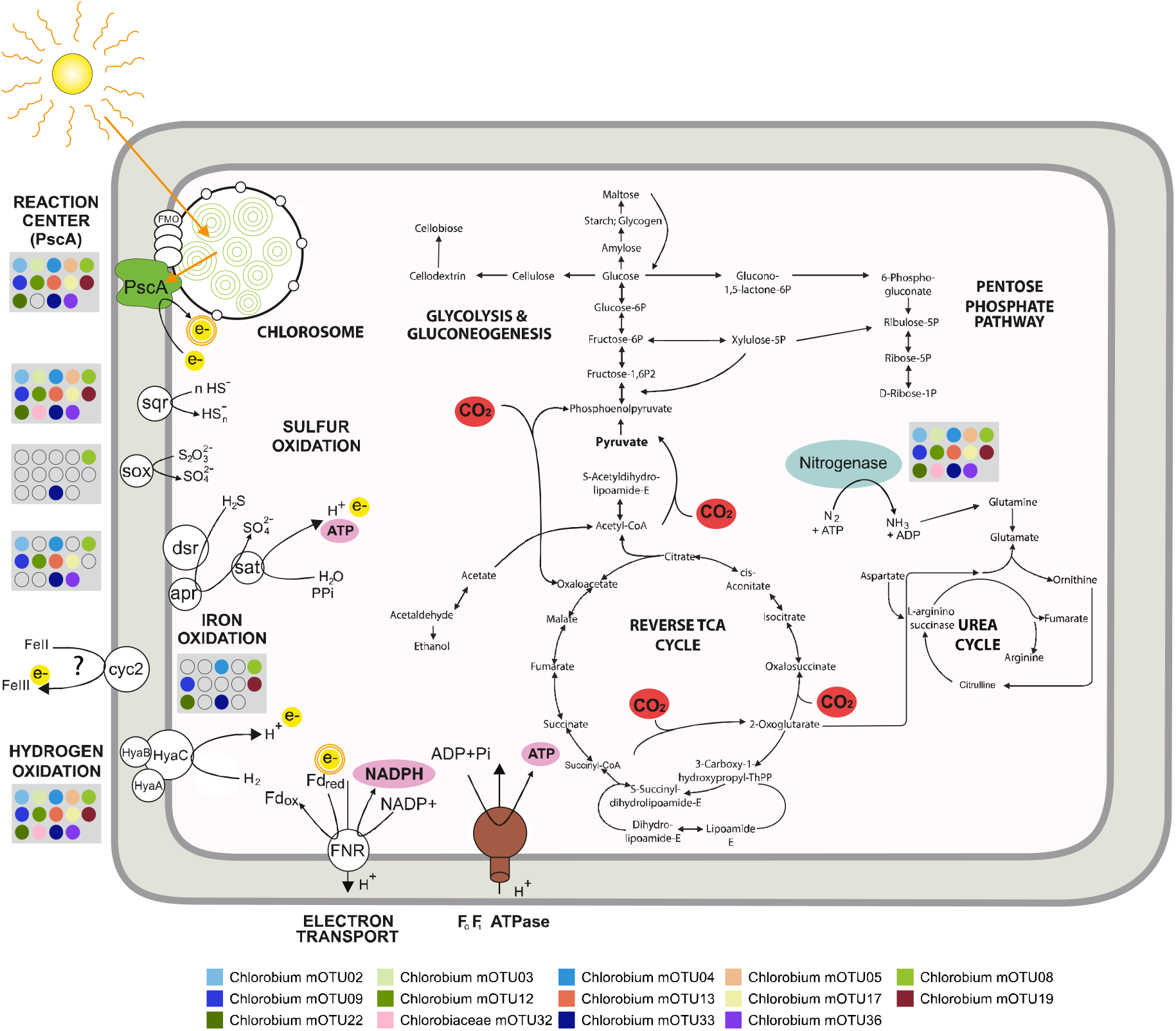
Metabolic potential of the 14 most abundant *Chlorobia* mOTUs found in the studied freshwaters, with focus on photosynthesis, electron donor, electron transport, and carbon fixation. Blue: the nitrogenase. Green: the chlorosome and photochemical reaction center PscA. Yellow: the electrons being donated through oxidation reactions and photosynthesis. Pink: reductant (NADPH) and chemical energy (ATP) produced during oxidation reactions and photosynthesis. Red: Pathways of carbon assimilation through the reverse tricarboxylic acid cycle, as well as anaplerotic gluconeogenesis steps, using electrons derived from inorganic compound oxidation and energy derived from photosynthesis. The photosystem uses electrons derived from sulfide, hydrogen, thiosulfate, iron oxidation and use light to activate them. This allows proton pumping and ferredoxin reduction. Ferredoxin reduction is linked to photosystem activity depicted with the double orange circle in electron. In the grey boxes there are circles where we colored according to presence/absence of the genes in the respective mOTUs in the legend. Gene product abbreviation, PscA – Photosystem I P700 chlorophyll a apoprotein A1, sqr – sulfide-quinone oxidoreductase, dsr – reverse dissimilatory sulfite reductase, apr – adenylylsulfate reductase, sat – sulfate adenylyltransferase, sox – thiosulfohydrolase, cyc2 – iron-oxidizing outer-membrane *c*-type cytochrome, Hya – group 1d [NiFe]-hydrogenase, FNR – fumarate and nitrate reductase, Fd – ferredoxin.

Depth stratification of *Chlorobia* mOTUs was evident when mapped reads were visualized as relative abundance of the total metagenomic reads per depth (Figure 4). As expected, our study showed that *Chlorobium* populations were mostly found below the oxycline (Figure 4) and some mOTUs were highly abundant. Most notably, for Lake Mekkojärvi (Finland), *Chlorobium* sequences comprised 57% of the community by relative abundance at some depths (Figure 4G). These high levels are in line with previous studies showing that this class can constitute 12 – 47% of reads in lake ecosystems (30, 31, 37, 38). Our data shows one clear case in which multiple *Chlorobium* populations typically co-inhabit the same lake with spatially separated niches. Specifically, mOTU02 and mOTU03 demonstrated differential depth preferences in August 2015 based on Lake Mekkojärvi samples (Figure 4G). Similar findings were previously reported in Trout Bog, where one of the populations was recovered only in the lowermost water layers (i.e., mOTU07, GSB-A/Chlorobium-111), whereas the other populations had broader distributions (e.g., mOTU58, GSB-B) (29, 31). It has been previously postulated that such niche specialization may reflect distinct pigment absorbance profiles (31, 39).

Several of the putative endemic mOTUs were temporally stable within their respective lakes. For example, we sampled Lake Lomtjärnen (Figure 4B) at four timepoints: March 2016 (under ice and snow cover), April 2016 (ice cover only), September 2017 (ice free) and July 2018 (ice free). The oxycline is lower in the water column when the lake is ice free and *Chlorobium* populations were abundant below the oxycline at all timepoints. The mOTUs residing in this lake were mostly endemic to the lake or to the region and they were consistently recovered at the different time points. It is also notable that the abundance of the different mOTUs changed according to the sampling time. For example, mOTU13 increased in relative abundance for the September sampling point. This allows us to speculate that, for endemic mOTUs, some may have specific seasonal niches. A more temporally resolved sampling effort would help test this hypothesis.

### The metabolic potential of the most abundant Chlorobium groups

We investigated the metabolic potential of all *Chlorobium*-associated mOTUs by inspecting the functionally annotated genes via eggNOG-mapper assigned K0 numbers. Combining the power of ANI clustering and statistical approaches helps calculate the probability that some genes are present in the core or accessory genomes of the “species”, despite the incompleteness of some of the MAGs and SAGs. In our data, 42 out of the 71 *Chlorobia* mOTUs were composed of more than one genome, and hence we find power in replication (40). For example, mOTU08 includes 34 MAGs with a completeness average of 95%, and the mOTU encodes most genes displayed in the core (Figure 2), except for HyaB, which is putatively within the accessory genome. Moreover, in order for a gene to be completely missing from the mOTU, the gene has to be absent in all of the MAGs and SAGs for the mOTU. Unlike traditional MAG-based studies, the absence of a gene in our mOTU data means that there is a high likelihood that the gene is really missing from the annotation or the population.

For individual mOTUs, we found an average of 710 annotated genes with designated KEGG functions. With these mOTUs, we were able to investigate the core metabolic functions of freshwater members of the genus *Chlorobium*. As expected, given the photolithoautotrophic lifestyle of cultured *Chlorobia*, core features of the genomes included all genes related to glycolysis and gluconeogenesis, reverse TCA cycle, and chlorophyll and bacteriochlorophyll biosynthesis (Figure 5; Figure S1A). These findings are consistent with previous studies showing very minor differences in the genes encoding the photosynthetic Type 1 reaction center unit in this class (41). We also observed molybdenum- and iron-nitrogenases widely encoded by the representatives of the class *Chlorobia* (Figure S1C). Thus, light harvesting, carbon fixation, and nitrogen fixation appear to be conserved core metabolic features of freshwater *Chlorobia* members.

In contrast to functions that were widely encoded, *Chlorobia* mOTUs significantly varied in their capacity to use electron donors (i.e., hydrogen, sulfide, thiosulfate, and iron). With respect to sulfur compounds, most mOTUs were capable of oxidizing sulfide via one or more enzymes. Sulfide-quinone oxidoreductases (Sqr) were encoded by all but two *Chlorobium* mOTUs (Figure 5 and S1B), whereas reverse dissimilatory sulfite reductases (i.e., DsrA) and flavocytochrome *c* sulfide dehydrogenase (*FCC*) were encoded by 71% and 53% of the mOTUs, respectively (Figure 2). A more restricted trait was the capacity to oxidize thiosulfate via thiosulfohydrolase (i.e., SoxB), which was only present in 14% of the *Chlorobium* mOTUs but in all *Chlorobaculum* mOTUs (Figure 2 and Table S2). Others have suggested that members of the Class *Chlorobia* have adapted to different environments by acquiring different electron transfer complexes through horizontal gene transfer (41–43). Even though genes for the Dsr and Sox complexes may have been horizontally acquired in this class (41), our phylogenetic trees show that DsrA and SoxB form monophyletic clades, consistent with a model that they were each acquired on one occasion during the evolution of *Chlorobia* and have potentially been lost from certain clades (Figure 2, Figure 6A and B).

**Figure 6.**
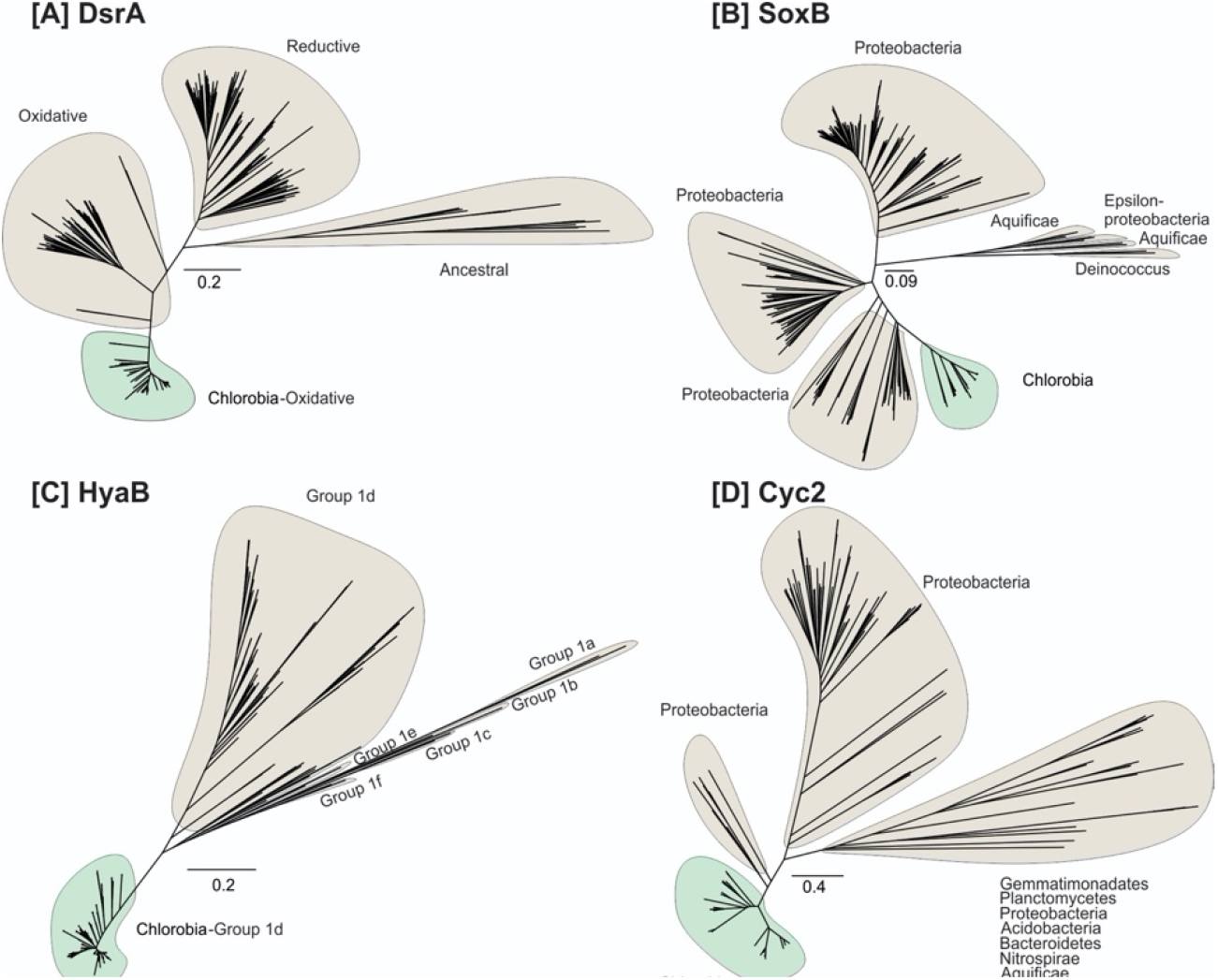
Maximum-likelihood phylogeny of the recovered protein sequences from reconstructed Chlorobia MAGs/SAGs (highlighted in green) together with the reference protein sequences recovered from NCBI GeneBank, with the exception of hydrogenases for which sequences were collected from HydDB (highlighted in gray). Panels show the phylogeny of (A) dissimilatory sulfite reductase, DsrA (n=551), (B) Thiosulfohydrolase, SoxB (n=294), (C) Group 1d NiFehydrogenase, HyaB (n=579), and (D) Iron oxidizing cytochrome, Cyc2 (n=306) protein sequences.

Genes encoding HyaB and HyhL (group 1d and group 3b [NiFe]-hydrogenases, respectively) were detected in 82% and 91% of the *Chlorobium* mOTUs, respectively (Figure 2, 6C and S1C). The group 1d enzymes are known to support hydrogenotrophic respiration and anoxygenic photosynthesis in diverse bacterial lineages (44–46). It is likely that the electrons liberated by these enzymes are transferred to the photosynthetic reaction center to reduce ferredoxin and NADP^+^ as a source of reductant for carbon fixation, nitrogen fixation, and other biosynthetic processes (47). Phylogenetic analysis confirmed that these subunits formed monophyletic radiations together with the hydrogenases retrieved from *Chlorobia* reference genomes (Figure 6C). The physiological role of group 3b hydrogenases (HyhL) in *Chlorobia* genomes is less clear. These bidirectional cytosolic enzymes are likely to support either hydrogenotrophic carbon fixation or facultative hydrogenogenic fermentation in this class (Figure S1D) (48, 49). Although uptake hydrogenases have previously been reported for *Chlorobium* members (50), the ecological significance of H_2_ metabolism has been overlooked in this class. Overall, the results suggest that H_2_ metabolism is an ancestral and conserved trait of the class *Chlorobia*. This is also consistent with the widespread distribution of [NiFe]-hydrogenases across bacterial phyla (51, 52) and the conservation of group 1d lineages in sister classes of the phylum *Bacteroidota*.

Our study also shows that 40% of the *Chlorobium* mOTUs harbor *cyc2* genes (Figure 2 and 6D), encoding a potential outer-membrane *c*-type cytochrome capable of iron oxidation (30, 53). This suggests that the distribution of the *cyc2* gene among *Chlorobium* genomes in lakes is higher than previously recognized. If the *cyc2* gene allows for extracellular electron transfer or ferrous iron oxidation, as speculated based on comparative genomics of cultured photoferrotrophs (30), this implies that *Chlorobia* members could be profoundly important for iron biogeochemistry in lake systems globally.

We did not find any mOTU encoding all the putative oxidation genes in the core genome (Figure 2 and 5). Metabolic flexibility is a key factor governing distributions of taxa across ecosystems (54). However, in this study we did not find a correlation between how widespread an mOTU is and their capacity to use different electron acceptors. A broader global sampling could reveal if metabolic versatility governs prevalence and abundance of *Chlorobium* bacteria globally.

### Outlook of ecological roles of Chlorobium

At a first glance, *Chlorobia* members appear to populate a very specific niche in water columns of global lakes and ponds. As we confirm in our datasets, they thrive under the oxycline in the anaerobic layer where light is still available. They have evolved physiological adaptations to low light intensities (3) and therefore seem to do particularly well in freshwater systems characterized by high concentrations of colored dissolved organic carbon (DOC) that absorb the incident solar radiation (55). In these lakes, solar radiation only penetrates to shallow depths in the water column, constraining populations of *Cyanobacteria* and most other photoautotrophs to the surface layer (56). *Chlorobium* members may contribute up to 83% of the total annual productivity in such humic waters (24) and also seem to play an important role as diazotrophs (15, 16). Despite the very specific niche of *Chlorobium*, our work illustrates considerable metabolic versatility with regards to oxidation genes that can be used to obtain electrons for carbon fixation. By their variable capacity to recycle sulfide, hydrogen, and iron produced by dissimilatory sulfate reducers, hydrogenogenic fermenters, and dissimilatory iron reducers, *Chlorobium* may potentially influence the entire microbiome of anoxic freshwaters (57). Our analyses describe the metabolic potential of environmentally relevant *Chlorobium* genomes along with their geographical distributions. Our results reveal both endemic and cosmopolitan clades but we were unable to link metabolic versatility to those distributions. Instead, the distributions of *Chlorobium* populations seem to distribute largely based on ecological factors. Our study also identified clades of *Chlorobium*, with abundant and cosmopolitan geographical distributions, that do not show a monophyletic clustering. In order to pinpoint the exact physicochemical background providing them with this ecological advantage, we need more comprehensive metadata and better geographical resolution. Nevertheless, our study highlights the ecological diversity of this enigmatic bacterial lineage.

## Materials and methods

### Collection of lakes and pond metagenomes

We obtained 265 metagenomes from 42 locations that spanned subarctic to tropical regions as a part of a project aiming to study microbial diversity in anoxic freshwater environments (Figure 1; Table S3) (28). Oxygen concentrations were measured with a YSI 55 oxygen probe (Yellow Springs Instruments, Yellow Springs, OH, USA). Moreover, we added six metagenomic libraries generated for a *Chlorobium* study in Canada focusing on iron oxidation (30) and three more metagenomic libraries are part of a 15-year timeseries study of Trout Bog (29).

### Genome collection

Assembling and binning the 265 metagenomes resulted in ~12,000 metagenome-assembled genomes (MAGs) from water bodies in 5 different countries (28). Of these, 454 belonged to the class *Chlorobia* and were used in this study. In brief, the raw data was trimmed using Trimmomatic (version 0.36) (58). The trimmed data were assembled using MEGAHIT (version 1.1.13)(59) with default settings. Two types of assemblies were performed: single sample assemblies for all samples individually and 53 coassemblies that were performed mostly on lake-specific metagenome data. Detailed descriptions of the sample collection, metadata, sample processing, metagenome generation and analyses methods are reported elsewhere (28).

The relevant quality controlled reads were mapped to all the assemblies using BBmap (60) with default settings and the mapping results were used to bin the contigs using MetaBAT (version 2.12.1, parameters --maxP 93 --minS 50 -m 1500 -s 10000) (61). Moreover, we collected genomes of the class *Chlorobia* from the GTDB and several other published MAGs that fulfilled the medium quality threshold of ≥50% completeness and ≤5% contamination (62). In total, we compiled 509 genomes that included 454 MAGs and 19 SAGs that were from stratified water bodies in Sweden, Finland, Canada, Switzerland, and Puerto Rico (28), seven MAGs from enrichment culture or lake metagenomes of lakes at the International Institute for Sustainable Development Experimental Lakes Area (IISD-ELA; near Kenora, Canada) (30), four MAGs from Trout Bog (29, 31), and 25 genomes (MAGs, SAGs or isolates) from the GTDB (2, 63). Completeness and contamination was assessed with CheckM (31).

### Single cell collection

In brief, sorting was done in 2016 on a MoFlo Astrios EQ sorter (Beckman Coulter, USA) using a 488 and 532 nm laser for excitation, a 70-μm nozzle, a sheath pressure of 60 psi and 0.1 μm filtered 1 x PBS as sheath fluid. An ND filter ND=1 and the masks M1 and M2 were used. The trigger channel was set to forward scatter (FSC) at a threshold of 0.025% and sort regions were defined on autofluorescence using a 532 nm laser and band pass filters 710/45 and 664/22. Sorted plates were stored at −80°C. The whole genome amplification was run using the REPLI-g Single Cell Kit (QIAGEN) following the manual’s instruction but with reduced total reaction volume to 12.5 μl. Amplified DNA was mixed thoroughly by pipetting up and down. The DNA was screened for bacterial 16S rRNA genes and DNA of confirmed *Chlorobia* members was sent for sequencing as previously described (28).

### mOTU analysis

Average nucleotide identity (ANI) for all genome pairs was computed with FastANI (version 1.3) (64) and the genomes were then clustered into metagenomic operational taxonomic units (mOTUs) with 70% completeness and 5% contamination thresholds. Genome pairs with ANI values above 95% were clustered into connected components. Additionally, less complete genomes (above 50% yet below 70% completeness) were recruited to the mOTU if its ANI similarity was above 95%. The classification of the genomes was done using GTDB-Tk and sourmash (version 1.0) (64, 66). The most complete MAGs per mOTU were selected as representatives and a phylogenetic tree was calculated using GTDB-Tk (version 102 with database release 89) with *Chloroherpetonaceae* as an outgroup (63). The aligned genome tree was loaded and curated in iTOL (version 5.5.1) (65). Moreover, functions (as in KO numbers) were classified as core if the probability of their presence-absence profile is higher under the assumption that it is in every genome, assuming their incompleteness, then the converse probability as computed by mOTUpan (https://github.com/moritzbuck/mOTUlizer).

### Abundance profiles

To explore abundance profiles, metagenomic libraries were subsampled to 1,000,000 reads (Table S2). Metagenomic reads were then mapped to all collected *Chlorobia*-associated genomes (Table S1). Reads were mapped with 100% identity using BBMap (60). The results were then normalized to relative abundance of mOTU per metagenome. All relative abundances of all samples in each location were averaged. The cutoff for presence of a mOTU was set to 0.03% read abundance per site. Heatmaps of abundance (Figure 3) were calculated and plotted using the R packages ggplot2 and phyloseq with parameters NMDS and Bray-Curtis for choosing the order of the x-axes (66–69). Depth-discrete abundance profiles were plotted using R packages ggplot2, phyloseq, and cowplot (67–69).

### Metabolic genes

The metabolic potential of the *Chlorobia* MAGs, SAGs, and reference genomes were reconstructed based on eggNOG-mapper (70, 71) annotations. Ten metabolic marker gene products (PsaA – Photosystem I P700 chlorophyll a apoprotein A1, AclB – ATP-citrate lyase beta-subunit, NifH – nitrogenase iron protein, FCC – flavocytochrome *c* sulfide dehydrogenase, Sqr – sulfide-quinone oxidoreductase, DsrA – dissimilatory sulfite reductase A subunit, SoxB – thiosulfohydrolase, Cyc2 – outer-membrane monoheme *c*-type cytochrome, HyaB – group 1d [NiFe]-hydrogenase large subunit, HyhL – group 3b [NiFe]- hydrogenase) were annotated in reconstructed MAGs, SAGs, and reference genomes using Diamond (version 0.9.31) (72) against custom build databases for each marker protein (query coverage threshold of 80%). A 50% identity threshold was used for all marker gene products except for an 80% threshold used for the PsaA protein. Annotations were further validated by reconstructing phylogeny.

### Single Gene phylogeny

Maximum-likelihood trees were constructed using the amino acid sequences of six metabolic marker proteins (PsaA – Photosystem I P700 chlorophyll a apoprotein A1, NifH – nitrogenase iron protein, Sqr – sulfide-quinone oxidoreductase, DsrA – sulfite reductase A subunit, SoxB – thiosulfohydrolase, Cyc2 – outer-membrane monoheme c-type cytochrome, HyaB/HyaA – group 1d [NiFe]-hydrogenase large subunit and small subunit, HyhL – group 3b [NiFe]-hydrogenase large subunit). Reference genes for each marker were collected from NCBI GeneBank, with the exception of hydrogenases for which sequences were collected from HydDB (46). For each metabolic marker protein, sequences retrieved from the *Chlorobia* mOTUs were aligned against reference protein sequences using ClustalW in MEGA7 (73, 74). Evolutionary relationships were visualized by constructing a maximum-likelihood phylogenetic tree. Specifically, initial trees for the heuristic search were obtained automatically by applying Neighbour-Join and BioNJ algorithms to a matrix of pairwise distances estimated using a JTT model, and then selecting the topology with superior log likelihood value. All residues were used and trees were bootstrapped with 50 replicates.

### Data availability

All datasets are available in public repositories with NCBI project accession numbers PRJEB3868, PRJNA518727, and PRJNA534305. Accession numbers of the genomes are available in Table S1, and the accession numbers of the metagenomes are available in Table S3. Assembling and binning of the original dataset has been done with the scripts available at https://github.com/moritzbuck/metasssnake General processing scripts for this project are available at https://github.com/moritzbuck/0026_Chlorobi

## Acknowledgements

The work was primarily funded by Science for Life Laboratory, the Olsson-Borgh, Knut and Alice Wallenberg Foundations (grant KAW 2013.0091), and Kungl. Vetenskapsakademiens stiftelser (BS2018-0108). K.D.M. acknowledges funding from the United States National Science Foundation Microbial Observatories program (MCB-0702395), the Long-Term Ecological Research Program (NTL-LTER DEB-1440297), and an INSPIRE award (DEB-1344254). J.D.N. acknowledges Discovery Grant and Strategic Partnership Grant for Projects funding from the National Sciences and Engineering Research Council of Canada (NSERC). S.B acknowledges funding from the Swedish Research Council and the Swedish Research Council Formas. The authors acknowledge additional support and resources from the National Genomics Infrastructure in Stockholm funded by Science for Life Laboratory, the Swedish Research Council, and SNIC/Uppsala Multidisciplinary Center for Advanced Computational Science for assistance with massively parallel sequencing and access to the UPPMAX computational infrastructure. The funders had no role in study design, data collection and interpretation, or the decision to submit the work for publication

SLG and SP conceptualized the research idea. All authors provided data. MB, MM, SLG, JMT, CG and KDM performed data analysis. SLG, MM, CG and SP did literature searches. SLG, MM, CG and SP drafted the manuscript and all authors contributed to writing and editing of the manuscript.

