## Supplemental Figure S1 for "Freshwater *Chlorobia* exhibit metabolic specialization among cosmopolitan and endemic populations"

**Figures and additional files**

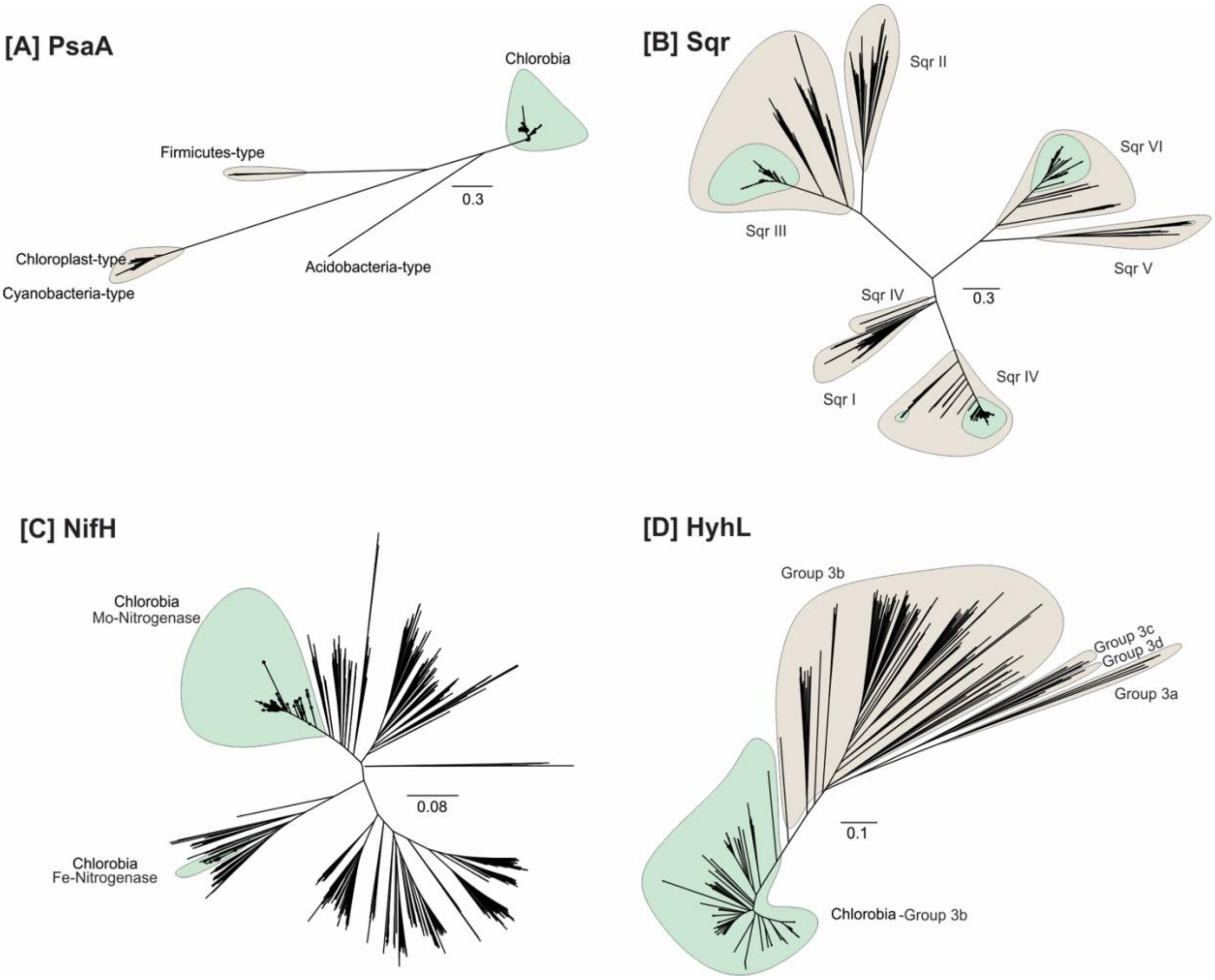


**Figure S1**. Maximum-likelihood phylogeny of the recovered protein sequences from reconstructed Chlorobia MAGs/SAGs (highlighted in green) together with the reference protein sequences recovered from NCBI GeneBank, with the exception of hydrogenases for which sequences were collected from HydDB (highlighted in gray). Panels show the phylogeny of (A) Type I reaction center subunits, PsaA (n=472), (B) sulfide quinone oxidoreductase, Sqr (n=1318), (C) Nitrogenase, NifH (n=1895) and (D) Group 3b NiFe hydrogenase HyhL (n=583) protein sequences.

**Table S1**. Accession number and statistics of all genomes in this study.

**Table S2.** mOTU statistics. In the gene content >0 means present in core, <0 means present in auxiliary and -10 absent completely.

**Table S3**. Metagenome metadata and accession number.
